# Territorial behavior as a route of social microbial transmission in an asocial mammal

**DOI:** 10.1101/2024.10.28.620674

**Authors:** Lauren Petrullo, Quinn Webber, Aura Raulo, Stan Boutin, Jeffrey E. Lane, Andrew G. McAdam, Ben Dantzer

## Abstract

Microbial transmission is a major benefit of sociality, facilitated by affiliative behaviors such as grooming and communal nesting in group-living animals. The spread of microbial symbionts through these pathways, and their incorporation into host microbiomes, can enhance host health and fitness by contributing to pathogen protection and metabolic flexibility. Are pathways that facilitate microbial transfer across hosts also present in animals that do not form social groups because territoriality limits social interactions and prevents group formation? Here, we addressed this question by combining longitudinal sampling of individual gut microbial communities, demographic data, and dynamic behavioral and spatial measures of territoriality from a non-social, highly territorial small mammal: wild North American red squirrels (*Tamiasciurus hudsonicus).* As squirrel densities increased, individual gut microbial communities became richer and more phylogenetically diverse, while among-individual differences in composition decreased. This pattern was characterized primarily by increases in obligately anaerobic and non-sporulating taxa with little to no tolerance for oxygen-rich environments, suggesting social rather than environmental routes of transmission. Moreover, territorial intrusions—in which conspecifics were found on within an individual’s territorial space—increased gut microbial diversity among individuals defending larger territorial spaces. Using an intrusion-based social network analysis, we found that that pairs with stronger social association (via intrusions) exhibited higher gut microbial similarity. Taken together, our findings provide some of the first evidence for social microbial transmission in a non-social species, and suggest that increased density and territorial behavior can diversify and homogenize host gut microbial communities despite social isolation.

## INTRODUCTION

The benefits of sociality are widespread. Prosocial behaviors such as grooming and communal nesting can reduce stress [1, 2], promote social cohesion [3], and improve access to food [4–6] and mates [7, 8]. Among these benefits is the host-to-host transmission of symbiotic microbiota (i.e., social microbial transmission), which despite an historical underappreciation has recently garnered substantial interdisciplinary interest [9]. At the individual level, social microbial transmission can enhance pathogen protection [10, 11] and support microbial recovery following ecosystem perturbations (e.g., after antibiotic administration [12]). More broadly, the social transmission of commensal microbiota can buffer against stochastic extinction events caused by bottlenecks over evolutionary time, preserving microbial diversity within a population [13]. While deterministic mechanisms (e.g., genetic effects and parent-offspring transmission) undoubtedly shape the host microbiota [14, 15], social pathways of microbial transfer can explain as much, or more, of host-to-host microbial similarity compared to relatedness, at least in social species [16–18]. Together, these patterns point to a central role for social microbial transmission in how animals acquire, maintain, and potentially fine-tune their microbiota in response to changing environmental and physiological demands [19, 20].

Social isolation–even when transient–can sever microbial transmission pathways, with potentially detrimental effects for host health and fitness [21]. As the vast majority of prior work has focused on highly social species, particularly those living in social groups (e.g., non-human primates and voles, [22–24]), whether such pathways exist in asocial species is unknown [23]. Nonetheless, non-social systems represent valuable models for uncovering the socio-ecological boundaries of host-to-host microbial transmission, and for disentangling the relative roles of vertical, social, and environmental routes of host microbial acquisition. Expanding our understanding of social microbial transmission to non-social systems is particularly timely as growing threats of natural disasters [25], pandemics [26], and habitat fragmentation [27] threaten to disrupt social connectedness, and thus the social transfer of beneficial microbiota.

Here, we test the hypothesis that fluctuations in conspecific density and territorial behavior in a non-social small mammal, the North American red squirrel (*Tamiasciurus hudsonicus)*, can facilitate social microbial transmission. In the southwest Yukon, red squirrel population densities fluctuate dramatically year-to-year as a result of the boom-bust dynamics of their primary food source, white spruce (*Picea glauca*, [28]). Food booms (mast years) occur episodically every 4-7 years, driving two major characteristics of our system: the coupling of food availability and squirrel density in space and time, and the larder hoarding of spruce cones by individual squirrels [29]. Hoards are centrally located within individual, stable, and non-overlapping territories that squirrels vigorously defend from intruders [30, 31]. However, territorial phenotypes (e.g., the size of an individual’s territory plus their ranging space, hereafter “territorial space”, and conspecific intrusions into this space) vary within and among individuals as a function of fluctuating population density [32]. For example, although squirrel densities peak following a food boom, territorial intrusions typically decrease [32]. By contrast, in non-mast years, densities decrease as a result of food scarcity and intrusions become more frequent [32]. These fluctuations in food availability, population density, and territorial intrusions generate appreciable intra- and inter-individual variation in expected rates of social microbial transmission, offering a novel contrast to the more static measures of sociality (e.g., bonded social relationships, social rank) commonly explored in similar studies on social animals [33].

In territorial animals, increased population densities and higher rates of territorial intrusions may directly (through incidental or aggressive physical contact) or indirectly (through environmental shedding via defecation and/or coprophagy) facilitate the transmission of microbiota between otherwise non-interacting hosts. We predict that when squirrel densities are high, or territorial intrusions become more frequent, individuals will exhibit (1) greater gut microbial diversity, suggesting host incorporation of conspecific microbiota, and (2) weaker individuality, indicating the homogenization of microbial communities across hosts through social transfer [34]. Recent work in wild mice has demonstrated that signatures of social transmission are characterized by the spread of oxygen-intolerant microbiota [23, 35]. We therefore additionally predict that (3) these effects will be driven primarily by obligately anaerobic, non-sporulating bacteria with little to no capacity for survival outside of the host gut.

## MATERIALS AND METHODS

### Ethics statement

All animal care and use permits and approvals were granted by the University of Michigan Institutional Animal Care and Use Committee (PRO00009804, PRO00007805 and PRO00005866) and were in accordance with the Canadian Council on Animal Care Guidelines and Policies. All relevant international and institutional guidelines for the use of animals were followed. Fieldwork was permitted under Yukon Territorial Government Wildlife Research Permits (205 and 218) and Scientists & Explorer’s Permits (17–13S&E and 18-08S&E).

### Study site and population

The Kluane Red Squirrel Project (KRSP) is a long-term ecological field project that has been continuously monitoring North American red squirrels (*Tamiasciurus hudsonicus*) in the Kluane region of the southwest Yukon, Canada, since 1989 [29]. All data presented here were collected from 2008-2017 (except 2013) from 52 squirrels (41 females, 11 males) living on one of two control/unmanipulated ∼40 ha study areas (“Kloo” or KL, “Sulphur” or SU) experiencing similar environmental conditions [60]. Here, we regularly follow all tagged squirrels from birth to death, identifying individuals by their uniquely labeled metal ear tags. Long-term monitoring included twice a year censuses of the entire population in May and September for assessment of territory ownership and squirrel population densities [29].

### Squirrel population density

We measured squirrel population densities by counting the number of squirrels defending individual territories at two scales. First, we calculated “grid-wise” densities based on the number of squirrels enumerated during the spring census divided by 40Lha (i.e. the area of each grid) for both KL and SU. This is a coarse measure of the overall squirrel density in an entire study area. Second, we then calculated local (i.e., neighborhood) densities as the number of squirrels defending individual territories within 50 m, 100 m, and 150 m of the focal individual’s territory centroid [48, 61]. Local density serves as a finer scale measure of squirrel density within an individual’s immediate area.

### Territorial phenotype

To generate territorial phenotype measures for individual squirrels, we followed previously published protocols [32]. Briefly, during the May census in each year, we recorded the location of each individual’s primary cache of spruce cones (i.e., midden) which is typically located near the center of its territory. Because squirrels typically defend the same territory for their entire life, and rarely change territories when new ones become available (with the exception of females occasionally bequeathing their territory to one of their offspring [62–64]), we used this midden location as an anchor for the spatial location of each individual’s territorial space in each year [65]. We then measured territorial intrusions as the observation (via trapping or behavioral observations) of conspecifics on this space. More details in Webber et al., (2023) and below.

#### Defining territories

To determine territorial spaces of individual squirrels, we followed procedures outlined in Webber et al., (2023). Briefly, we used a combination of trapping and behavioral observations to generate estimates of territorial space use for each squirrel, and followed a series of conservative data inclusion criteria and validations to reduce bias associated with estimating territories. First, we removed all observations of squirrels that occurred outside of our study area (and thus lacked precise spatial locations). Second, we generated territorial space measures only for squirrels with at least 20 behavioral observations and/or trapping events in each year (i.e., between March 15 and September 1). For squirrels with >30 observations/day, we randomly picked 30 observations to include in the analysis (∼95% of squirrels had < 30 observations/day). Using these data, we generated 30% kernels as a conservative measure of the boundaries of an individual’s territorial space [32]. Despite this conservative analytical approach, we chose to define these estimates as territorial space rather than territories to be even more conservative in the event that our estimates capture both individually defended territories and potential ranging space used by a squirrel for breeding. Thus, we consider territorial spaces as an individually variable area around a squirrel’s strictly defended midden.

#### Territorial intrusions

We defined territorial intrusions as instances in which a squirrel was observed or trapped within the territorial space (as defined above) of a conspecific (hereafter, “owner”), excluding juveniles captured within the territorial spaces of their mothers in the first year [48]. Individual territories are stable such that squirrels with established territories rarely bequeath them [63], and any abandoned territories (e.g., due to death of the owner) are typically taken over by new individuals quite rapidly [31, 59]. Thus, we used a hierarchical temporal moving window approach to remove potential false positives in our measurement of intrusion events, described elsewhere [32]. In brief, we considered the tenure of territorial space ownership and the earliest and latest days of our trapping and behavioral observations for a particular squirrel each year. We then conservatively removed any intrusion events in which a squirrel was observed in the territorial space of an owner prior to the earliest trapping/observation day that year and after that owner’s last trapping/observation day that year if the owner did not own that territorial space in the following year.

### Fecal sample collection

We longitudinally sampled the 52 individuals in this study, collected 562 fecal samples (mean = 11 samples/individual, range = 1-34) resulting in 110 unique individual-grid-year combinations. Fecal samples were collected non-invasively and opportunistically from below traps during routine live-trapping events as previously described [29]. Fresh fecal samples were immediately stored on wet ice in the field until they were frozen at −20°C within 5 h of collection and kept at - 20°C before being shipped to the University of Michigan where they were stored at −80°C until analysis.

### 16S rRNA gene sequencing

We used 16s rRNA amplicon sequencing to determine the composition of the gut microbiota from fecal samples. Briefly, microbial DNA was extracted using the Qiagen MagAttract PowerMicrobiome kit (Qiagen, USA) following the manufacturer’s instructions, and extracted DNA was quantified using the Quant-iT PicoGreen dsDNA assay kit. The hypervariable V4 region of the 16S rRNA bacterial gene was then amplified using a dual-indexing approach to allow for sample assignment after multiplexing [66]. To amplify the V4 region, we used the following primers (forward: GTGCCAGCMGCCGCGGTAA, reverse: GGACTACHVGGGTWTCTAAT, [66]). Following amplification and indexing, libraries were pooled, spiked with PhiX to increase library complexity, and sequenced in two separate runs on an IlluminaMiseq using a MiSeq Reagent Kit V2 (500 cycles) at the University of Michigan Microbiome Core. Each run included 3 positive controls (ZymoBIOMICS Mock Community II) and at least 3 negative controls (H_2_O).

### Bioinformatics

We processed sequencing data using Quantitative Insights Into Microbial Ecology 2 (QIIME2; qiime2-amplicon-2023.9, [67]). We first demultiplexed raw reads (demux--) before running the Divisive Amplicon Denoising Algorithm 2 (DADA2) pipeline to collapse sequences into amplicon sequence variants (ASVs), which vary by a single nucleotide [36]. We denoised each sequencing run separately before merging to account for potential batch effects of separate runs [36]. After visualization of read quality, we trimmed reads at 240 bp to remove low-quality end portions of sequences. After filtering, trimming, merging, and chimera removal, we retained a total of 13,097,568 reads across 562 fecal samples (mean = 23,305 reads, SD = 8,595, range = 1,243 - 53,686). Negative control samples (H_2_O) had read depths ranging from 0 to 15, suggesting minimal kit contamination. We assigned taxonomy to all reads using the q2-feature classifier in QIIME2 against the most recent version of the SILVA database (SILVA 138 SSU Ref NR 99) based on 100% similarity to the reference sequence [68]. Assignment was universal at the phylum level (100%), and high at both the family (98%) and genus (97%) levels.

### Statistical analysis

We imported QIIME2 data into R (version 4.3.1, [69]) using *qiime2R* [70]. All models controlled for sample read depth (library size), either as a fixed factor or as a log-transformed offset when the dependent variable was raw read counts of microbial taxa. This allowed us to control for differences in sequencing depth among samples without subsampling, and therefore avoid the associated inflation in false positive rates and removal of meaningful sequence information associated with rarefaction ([71], but see [72]).

#### Calculation of among-individual microbial diversity (beta-diversity)

We quantified among-sample microbial diversity by calculating Jaccard similarity, which is an integrated and bounded measure of ASV presence/absence that does not rely on relative abundance in its calculation. We chose this metric because it is more sensitive than other similar beta-diversity metrics (e.g., Bray-Curtis, UniFrac) to signals of social microbial transmission [17, 35]. We first computed a Jaccard distance matrix in *phyloseq* [73] and visualized sample spread using Principal Coordinates Analysis (PCoA) in *vegan* [74]. We then performed marginal Permutational Multivariate Analysis of Variance (PERMANOVA) with 999 permutations to test for associations between gut microbiome composition and variables of interest using adonis2(). We controlled for individual ID in these models by setting ID as a blocking factor prior to running.

Tests for homogeneity of multivariate dispersion can be used to assess categorical group-level differences in beta-diversity and determine differences in the inter-individual variation in microbial community composition among groups. To determine whether dispersion differed at different population densities, we first binned all samples into low, medium, or high-density categories based on an even three-way split. We then used the *betadisper*() function, which uses a multivariate analogue of Levene’s test for homogeneity of variances, to calculate the relative distances of all samples from their group centroid in multivariate space (Jaccard distance). We extracted distances and ran a generalized linear mixed-effects model (family = beta regression) to test whether squirrel density (low, medium, high) predicted sample distance to centroid (dependent variable), controlling for grid, season, sample read depth, year, and individual ID (random effect).

#### Calculation of within-individual microbial diversity (alpha-diversity)

For all samples, we calculated three different measures of within-sample alpha-diversity: observed richness (number of unique ASVs per sample), Shannon Index (ASV richness and evenness), and Faith’s phylogenetic diversity (diversity of phylogenetic relationships among ASVs within a sample) using *phyloseq* [73] and *picante* [75]. Shannon indices were tukey-transformed to achieve residual normality prior to analysis.

#### Generation of predicted microbial functions

We used the Phylogenetic Investigation of Communities by Reconstruction of Observed States version 2 (PICRUSt2) pipeline to estimate microbial gene content and infer microbial function from amplicon sequencing data [37]. Briefly, ASVs were aligned to reference sequences using HMMER [76] and placed into a reference tree using EPA-NG [77] and gappa [78]. Data were normalized via castor [79] and mapped onto gene pathways using MinPath [80]. We investigated functional profiles based on Kyoto Encyclopedia of Genes and Genomes (KEGG) orthologs (KOs), which we then mapped onto general functional categories at Level 2 of the BRITE map [81]. One input sequence aligned poorly to the reference sequences and was therefore excluded from downstream analysis (ASV 2ce86713ae7cbabc639eb411191e6128). We assessed the accuracy of our predicted functional pathway data by calculating a weighted Nearest Sequence Taxon Index (NSTI) for each sample, which quantifies the extent of similarity between the ASVs within a sample and the reference genomes [82]. The mean NSTI score for our dataset was 0.11 ± 0.02, falling within the range of NSTI values obtained for mammalian gut microbiota in a validation of PICRUSt2 predictions against metagenomic data (mean NSTI 0.14 ± 0.06, [82, 83]).

#### Differential abundance of microbial taxa and functional pathways

Taxonomic data were agglomerated at the genus level, and transformed to relative abundance. We took a relatively liberal approach, using a sample-wise abundance threshold to retain microbial taxa with average relative abundance of at least 0.05% across all samples. To identify taxa whose abundances changed as a function of population density and territoriality, we constructed zero-inflated negative binomial mixed-effects models using *glmmTMB*. All models controlled for the effects of season, study area, and sequencing run (fixed factors), individual identity and year (random factors), and differences in sample library size by including log-transformed read depth as a model offset. For predicted functional data, we converted all functional pathway counts to relative abundance and retained all functional pathways at Level 2 of the BRITE map. To identify KEGG pathways that significantly differed in relative abundance with population densities, we ran a series of linear mixed-effects models including the same covariates and all identified pathways (N=43). For all models, we accounted for multiple comparisons and an inflated false discovery rate by adjusting all P values using the Benjamini-Hochberg correction [84].

#### Assignment of aerotolerance and sporulation capacity to microbial taxa

For the ten microbial genera found to be differentially abundant as a function of population density, we assigned phenotypes of facultative and obligate aerotolerance (anaerobic) and capacity for sporulation (sporulating, non-sporulating) using Bergey’s Manual of Systematics of Archaea and Bacteria [85]; where taxa were not available in Bergey’s Manual, we referred to Raulo et al. (2024) for assignment of microbial phenotype, particularly for more recently named and/or classified taxa [35]

#### Investigating the effect of territorial intrusions on gut microbial similarity using social network analysis and dyadic Bayesian regression

To test whether social behavior in the form of territorial intrusions predicted pairwise gut microbial composition, we first generated a social network based on intrusions between each pair using a lifespan overlap -corrected simple ratio index [86]. This index defined social association strength as the total number of unique intrusions between individuals A and B (# of times individual A invaded B, and B invaded A) divided by the total number of intrusions either A or B were part of (# of times individual A invaded B, and B invaded A + # of times A invaded anyone else or was invaded by anyone else + # of times B invaded anyone else or was invaded by anyone else). To generate this measure accurately we used a larger dataset of intrusions that included data from squirrels not fecal sampled for the current study [32].

We then used a dyadic Bayesian modeling approach to predict the level of microbiota sharing (Jaccard similarity) with social association strength among pairs of squirrels. This was made possible by a Dyadic Bayesian multi-membership beta regression model that allows random effect structures that can account for the inherent interdependence of pairwise data and the non-independence rising from repeated sampling of the same individual [17, 35]. Using the *brms* package [87], we constructed models with dyadic pairwise microbial similarity values (Jaccard similarity) as the response variable, excluding self-comparisons (e.g, pairwise comparisons of samples collected from the same individual) [17, 35]. All models used a Markov chain Monte Carlo sampler implemented with the *rstan* package. For the first model, we used the entire Jaccard distance matrix (N = 153,581 pairwise comparisons, N = 1,326 unique pairs) to determine how similarity in intrinsic (host age, sex), temporal (sampling season, year) and spatial (study area) variables predicted microbial similarity. We controlled for differences in read depth between the samples in a pair (fixed variable), as well as individual and sample ID by including a multi-membership random effect for ID (ID of individual A, ID of individual B of the pair) and sample (Sample A, Sample B of the pair). This model was run with 4 parallel chains threaded across 32 cores, each with 3,000 warm-up samples preceding 12,000 actual iterations. We used posterior checks to ensure reliable model performance, assessing chain convergence and ensuring that Rhat values were < 1.10.

For the second model we included social association as a main predictor and subsetted the data heavily to only include pairs from relevant temporal and spatial scales: Because pairs inhabiting different study areas rarely intrude upon each other’s territories, we subsetted the main distance matrix to only those pairwise comparisons capturing individuals inhabiting the same ∼40 ha study area. Because our measures of territorial phenotype were calculated annually for each individual, we further subsetted to only include pairwise comparisons between fecal samples collected in the same year, resulting in N = 25,381 pairwise comparisons across N = 436 unique pairs. We then ran the model predicting microbiota similarity with social association strength (the simple ratio index) controlling for covariates: season similarity same/different), sex similarity (same/different), age difference (numeric), read depth difference (numeric). We additionally controlled for potential spatial autocorrelation in microbiota (and thus environmental transmission) by including spatial distance between the centroids of both squirrels’ territories in this model. For both models, all continuous variables were scaled between 0 and 1 prior to analysis, unless they already naturally ranged between 0 and 1.

## RESULTS

### Socio-environmental structuring of the red squirrel gut microbiota

We profiled the gut microbiota of 52 individuals from 562 fecal samples using 16S sequencing resolved to amplicon sequence variants (ASVs, [36]). We detected 4,547 unique ASVs across all samples, with the majority of ASVs belonging to five major bacterial phyla. Overall, the red squirrel gut microbiota was dominated by bacteria within the phyla *Firmicutes* (mean relative abundance = 47.4%) and *Bacteroidota* (mean relative abundance *= 38.3%),* with fewer taxa assigned to *Campilobacterota, Proteobacteria, and Actinobacterota* (Figure 1A). Within these phyla, sixteen families were highly abundant (Figure 1B). The most abundant bacterial families were *Prevotellaceae, Muribaculaceae, Lachnospiraceae, Oscillospiraceae, Ruminococcaceae,* and *Bacteroidaceae,* each comprising on average between 9-15% of the total gut microbiota (Figure 1B).

**Figure 1.**
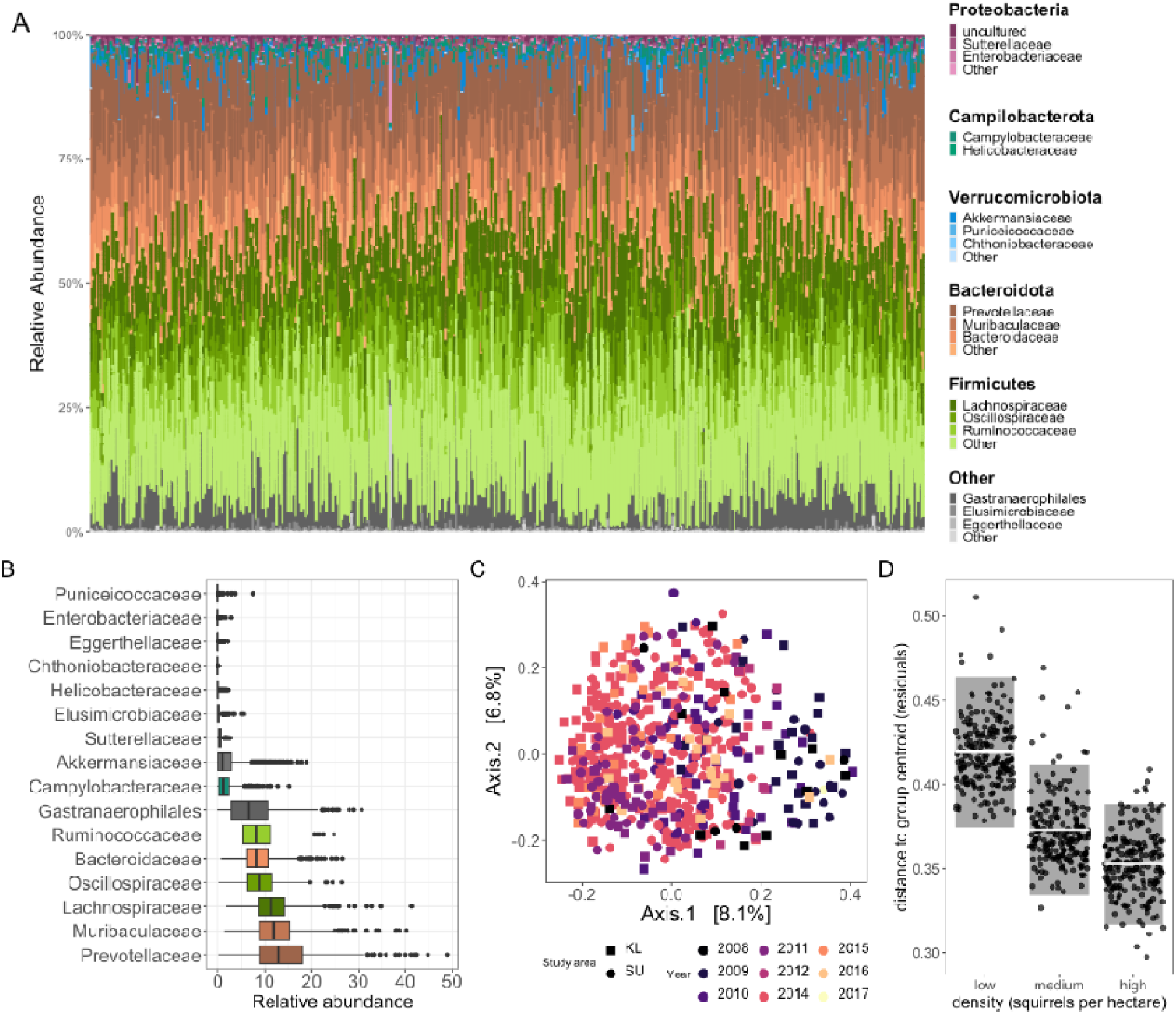
The red squirrel gut microbiota is shaped by temporal, spatial, and ecological factors. **(A)** Taxonomic composition of all samples, showing the five most abundant phyla and the three most abundant families within each phylum. **(B)** Variation in the relative abundance of the top microbial families (median relative abundance between 1-13%). (**C**) Spatio-temporal clustering by study area and year of sample based on Jaccard similarity indices (presence/absence). **(D)** Partial residuals from a generalized linear mixed-effects model (family = beta) testing for significant (*p* < 0.05) differences in individuality of the microbiota, measured as distance to the group centroid in Jaccard compositional space at different squirrel densities, controlling for individual identity.

Social, environmental, temporal, and host factors predicted variation in the red squirrel gut microbiota to different degrees. Host identity explained most of the variance in microbial community structure among samples (∼20%), indicating strong signatures of individuality within the microbiota (Table S1, Figure 1C). However, this individuality of the gut microbiota fluctuated as a function of the squirrel population densities. Interindividual differences in gut microbiota composition, measured as distances to group centroids (high, medium, and low density) in Jaccard compositional space, were smallest when densities were high and greatest when densities were low (estimate ± SE: −0.07 ± 0.02, z = −3.32, *p* = 0.0009; Figures 1D and S1, Table S3). After controlling for the effects of individual identity, temporal variables explained more of the variation in microbiota than spatial variables did, with year and sampling season explained 4% and 6% respectively. The study area within which a squirrel lived (∼40 ha space) explained just 0.9% of the total variance, and there was no effect of host age or sex on microbial composition (Table S2).

### More neighbors, more microbes

Across the 9 years of this study, squirrel densities fluctuated dramatically at both fine and coarse scales (Figures S2-S3), and there was appreciable individual variation in microbial richness as a function of density (Figure 2A). Squirrels with more neighbors (i.e., individuals defending territories) within 50 m of the center of their own territories exhibited greater gut microbial richness (estimate ± SE: 4.22 ± 1.96, t = 2.15, *p* = 0.03, Figure 2B), abundance-weighted diversity (Shannon Index, estimate ± SE: 53.0 ± 15.7, t = 3.38, *p* = 0.0008), and phylogenetic diversity (Faith’s Phylogenetic Diversity, estimate ± SE: 0.16 ± .008, t = 2.0, *p* = 0.04) than squirrels with fewer neighbors within 50 m (Table S4). As the radius within which we measured neighborhood density increased (e.g., to 100 m and then to 150 m), the relationship between squirrel density and ASV richness disappeared, but effects of density on abundance-weighted (Shannon Index) and phylogenetic diversity remained (Table S5). At the coarsest scale (i.e., total number of squirrels within an individual’s entire ∼40 ha study area), squirrel densities did not predict variation in individual gut microbial diversity by any alpha-diversity measure tested (Table S6).

**Figure 2.**
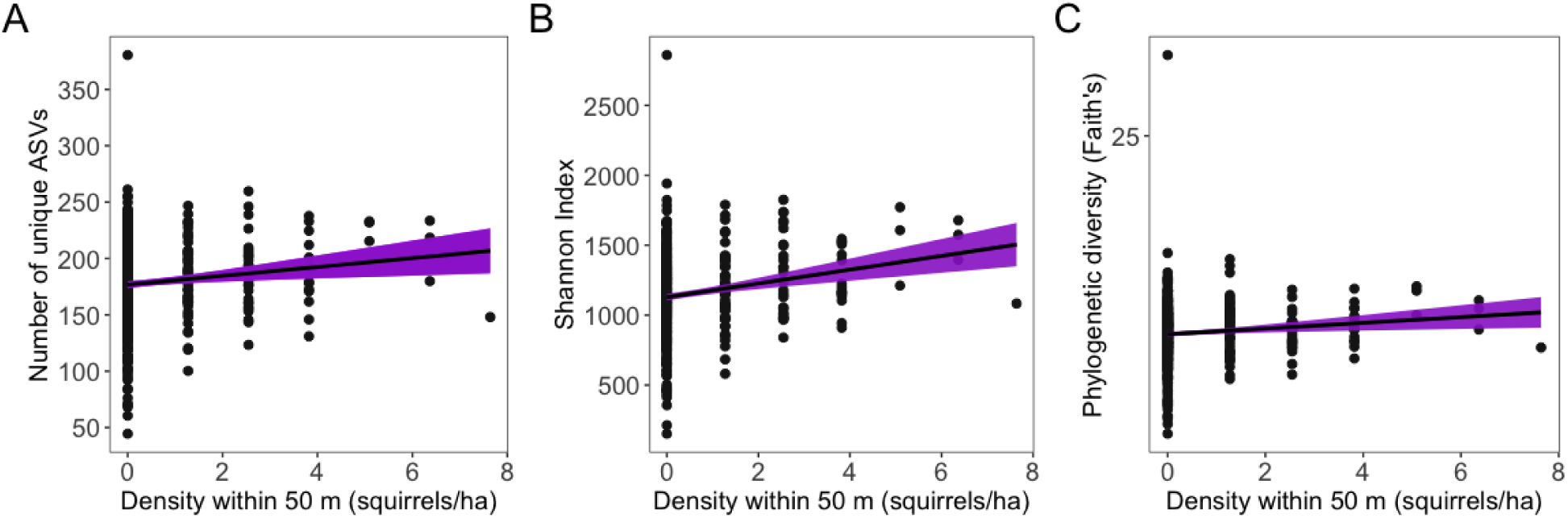
Elevated population density predicts greater gut microbial richness and phylogenetic diversity. At higher squirrel densities (within 50 m of the centroid of an squirrel’s territorial space), individual gut microbial communities exhibit significantly (*p* < 0.05) greater **(A)** ASV richess, **(B)** abundance weighted diversity (Shannon Index), and **(C)** phylogenetic diversity (Faith’s). Plots show results from linear mixed-effects models; points reflect samples (partial residuals); line and bands show fitted regression with 95% CIs.

To identify the microbial taxa driving the shift in community alpha-diversity with changes in population density, we modeled counts of bacterial genera as a function of squirrel density within 50 m of an individual’s territory, controlling for environmental, spatial, and host factors. As squirrel densities increased, genera in the family *Prevotellaceae* (including UCG-001 and Ga6A1-group) decreased in relative abundance (Figure 3A, Table S7). Several bacterial genera increased, including *Sanguibacter* (β = 0.527, *p_FDR_ =* 0.047)*, Herbinix* (β = 0.342, *p_FDR_ =* 0.026)*, Eubacterium_nodatum_group* (β = 0.276, *p_FDR_ <* 0.0001)*, Frisingicoccus* (β = 0.167, *p_FDR_ =* 0.047)*, Erysipelatoclostridium* (β = 0.132, *p_FDR_ =* 0.004)*, Gastranaerophilales* (β = 0.100, *p_FDR_ =* 0.035), and two genera in the family *Oscillospira* [NK4A215_group (β =0.089, *p_FDR_ =* 0.047) and UCG-005 (β = 0.084, *p_FDR_ =* 0.035), Figure 3A, Table S7]. Of the 8 microbial genera taxa that increased with increasing densities, five were obligate anaerobes while three were genera with at least some aerotolerance [i.e., facultative anaerobes, Figure 3A]. When densities were lower, two obligately anaerobic *Prevotella* [Ga6A1_group (β = −0.153, *p_FDR_ =* 0.040) and UCG-001 (β = −0.135, *p_FDR_ =* 0.011)] decreased in relative abundance. All ten microbial genera that varied with fluctuating squirrel densities were non-sporulating taxa, with the exception of two taxa for which sporulation capacity is unknown or unresolved.

**Figure 3.**
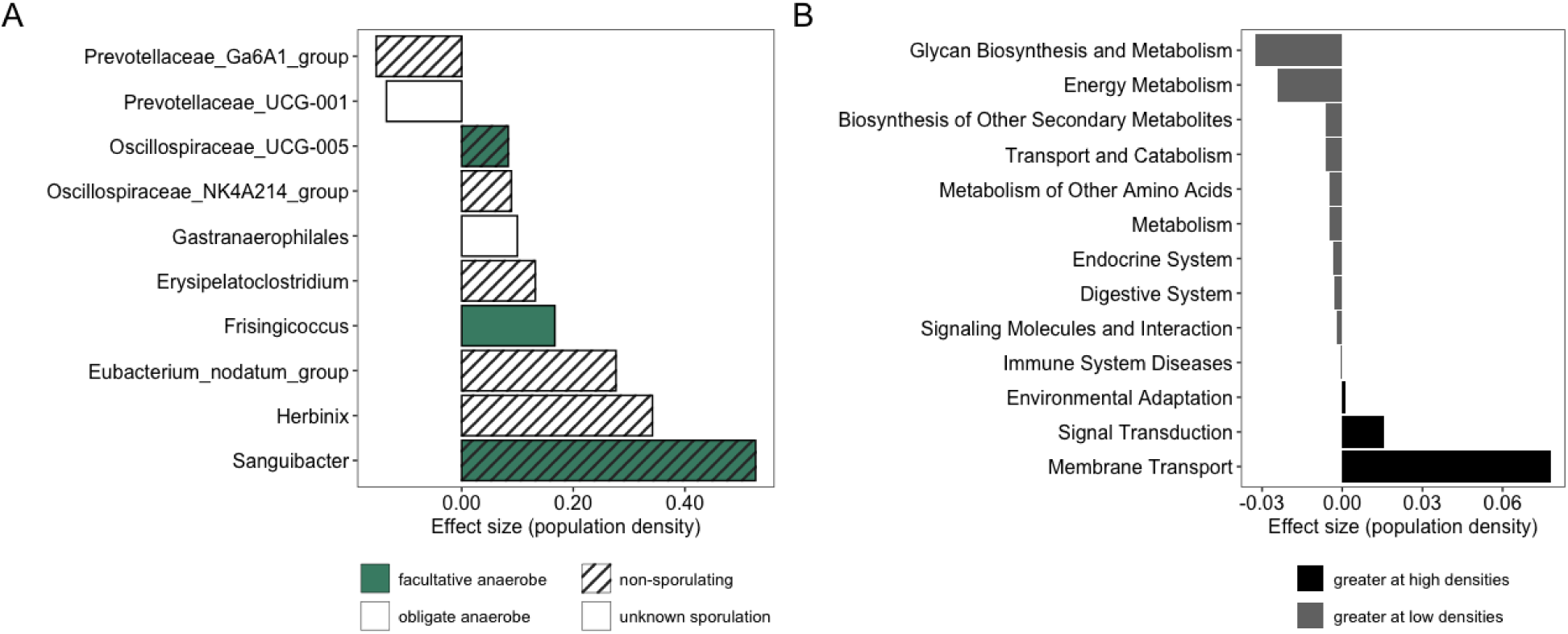
Shifts in the gut microbiota with changing density involve primarily anaerobic, non-sporulating bacteria and environmental sensing pathways. Density predicts differential enrichment and depletion of **(A)** microbial genera with variable levels of aerotolerance, and **(B)** microbial functional pathways. Plots depict results of mixed-effects models (A: zero-inflated negative binomial; B: linear) testing microbial genera (N = 84 genera) and predicted KEGG orthologs (N = 41) that significantly (*p*_FDR_ > 0.05) differ in relative abundance as a function of changing squirrel densities.

The functional capacity of the gut microbiota also changed with squirrel population density based on predicted metagenome function [37]. Overall, the red squirrel gut microbiota was dominated by functional microbial pathways related to membrane transport (median relative abundance = 10.4%, range = 8.2 - 12.7%), replication and repair (10.3%, 9.6 - 11.0%), and carbohydrate (9.7%, 9.3 - 10.8%) and amino acid (9.7%, 9.3-10.4%) metabolism (Figure S4). At higher squirrel densities, gut microbiota shifted towards a greater functional capacity for genes involved in environmental adaptation (β = 0.001, *p_FDR_ =* 0.046), signal transduction (β = 0.015, *p_FDR_ =* 0.008), and membrane transport (β = 0.078, *p_FDR_ =* 0.046, Figure 3B, Table S8). By contrast, at lower densities, functional pathways associated with major host organ systems and physiological processes such as energy metabolism (β = −0.024, *p_FDR_ =* 0.008), immunity (β = - 0.001, *p_FDR_ =* 0.008, digestion (β = −0.003, *p_FDR_ =* 0.003), and endocrine function (β = −0.003, *p_FDR_ =* 0.006) dominated (Figure 3B, Table S8).

### Territorial spaces and territorial intrusions interact to predict gut microbial diversity

The territorial space used by squirrels varied in size depending on host sex (males had larger territorial spaces, estimate ± SE: 0.27 ± 0.11, t = 2.33, *p* = 0.02) and neighborhood density within 50 m (higher densities predicted smaller territorial spaces, estimate ± SE: −0.02 ± 0.01, t = −2.01, *p =* 0.045, Table S9; [32]). There was no effect of neighborhood density within 100 m or 150 m on territorial space size. Individuals with larger territorial spaces also experienced a higher rate of territorial intrusions compared to those with smaller spaces (estimate ± SE: 0.06 ± 0.007, t = 9.58, *p* < 0.0001; Table S10, [32]).

The size of an individual’s territorial space was negatively correlated with gut microbial diversity, such that individuals with smaller territorial spaces exhibited greater species richness (estimate ± SE: −4.73 ± 2.16, t = 02.19, *p* = 0.03) and abundance-weighted diversity (estimate ± SE: - 40.07 ± 17.41, t = −2.30, *p* = 0.02) compared to those using larger spaces (Table S11).

Controlling for density, the frequency of territorial intrusions predicted gut microbial alpha-diversity, but only when interacting with territorial space size: for squirrels with larger territorial spaces, a higher rate of intrusions predicted greater microbial richness (estimate ± SE: 6.28 ± 2.03, t = 3.10, *p* = 0.002, Figure 4), abundance-weighted diversity (estimate ± SE: 41.07 ± 16.27, t = 2.53, *p* = 0.12) and phylogenetic diversity (estimate ± SE: 0.32 ± 0.08, t = 3.76, P = 0.0002; Table S11).

**Figure 4.**
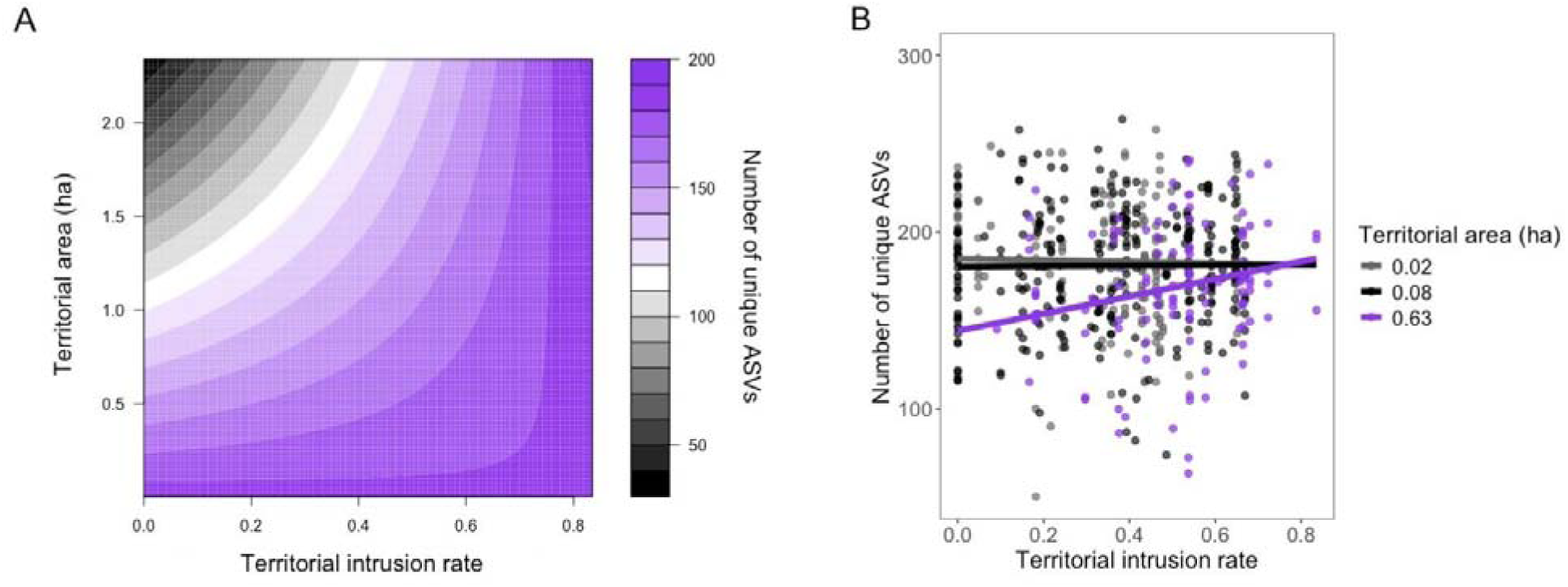
More frequent intrusions predict greater gut microbial diversity. (**A**) Surface plot showing slope and direction of the interaction between territory size and territorial intrusions on gut microbial richness. As territorial area (y axis) and the rate of territorial intrusions (x axis) increase, so does gut microbial richness (z). (**B)** Individuals with larger territorial spaces that experience more territorial intrusions exhibit greater gut microbial richness. Plot depicts partial residuals from a linear mixed-effects model controlling for density and other covariates (see Table S11), and three overlaid slopes at different territory sizes. Note one sample with microbial richness > 600 on the y-axis removed from the plot for visualization purposes but retained in the model.

### Territorial intrusions homogenize the gut microbiota of intruder pairs

The mean gut microbiota similarity (Jaccard similarity) across our entire dataset was 0.18 (se = 0.06), indicating that on average, pairs of squirrels shared approximately 18% of their gut microbiota. Using a Bayesian dyadic modeling approach (*brms*, [17, 35]), we investigated the pairwise variables that explained variation in the degree of this similarity across pairs of squirrels in our entire dataset. Broadly, pairs (N = 1,326 pairs) had more similar microbiota if they were sampled in the same year (posterior mean = 0.1, CI = 0.01 - 0.01) and season (posterior mean = 0.14, CI = 0.14 - 0.14), if they inhabited the same study area (posterior mean = 0.02, CI = 0.01 - 0.02), and if they were similar in age (posterior mean = −0.01, CI = −0.02 - 0.0, Table S12). There was also a strong effect of the difference in sample read depth (posterior mean = −0.86, CI = −0.88 - −0.85), but no effect of similarity in host sex on microbial similarity (Table S12).

To determine whether territorial intrusions facilitate social microbial transmission, we first generated an intrusion-based social association network for all squirrels in our dataset. Social association for each pair was calculated as the number of pairwise intrusions between squirrel A and squirrel B (the number A invaded B + the number B invaded A), divided by the total number of intrusions A and B were part of (Figure 5A). We then used a similar approach as above for dyadic pairwise comparisons, but limited comparisons to pairs inhabiting the same study area and sampled for microbiota in the same year (N = 436 unique pairs). Controlling for potential confounding effects of host age, sex, season, spatial proximity, and sample read depth, pairs with stronger social association (through territorial intrusions) had more similar gut microbiota, and this was the strongest effect in the model (after sample read depth; social association strength posterior mean = 0.22, CI = 0.14 - 0.29, Table S13, Figure 5B-C). Importantly, this effect of intrusion-based social association was independent of the effect of spatial distance on microbiota similarity (pairs in closer proximity had more similar gut microbiota regardless of any intrusions between them, posterior mean = −0.1, CI = 0.14 - 0.29, Table S13, Figure 5B).

**Figure 5.**
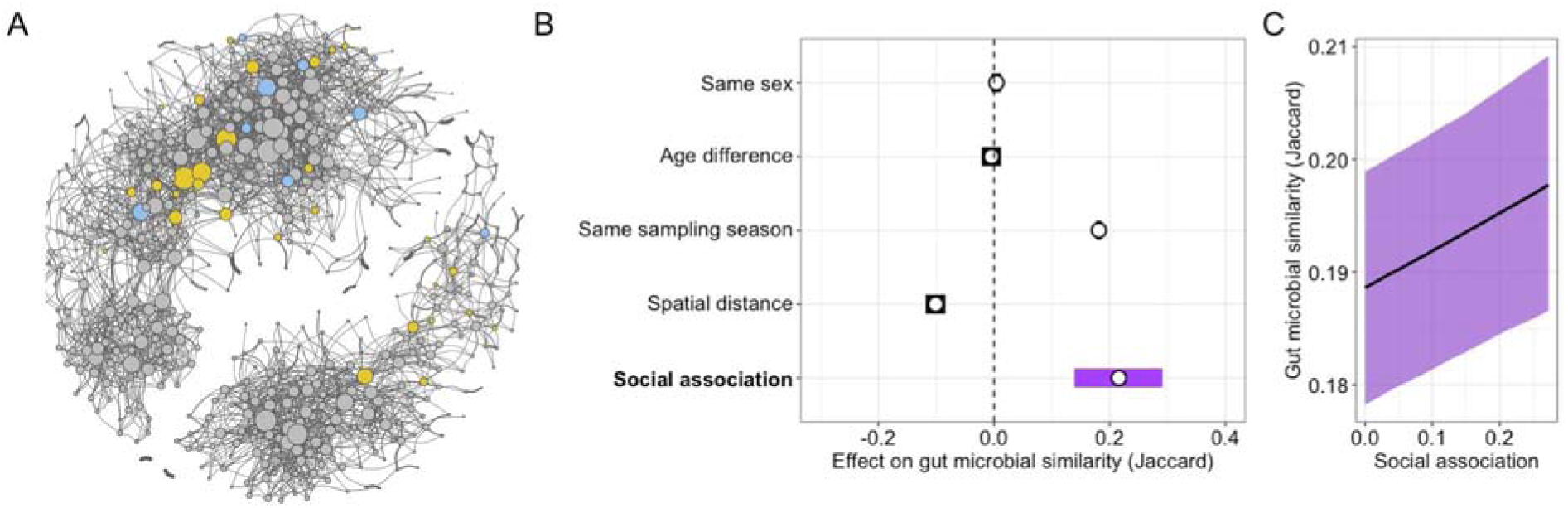
Dyads with stronger social association based on intrusions have more similar gut microbiota. **(A)** Social network based on social association index. Node size reflects the number of contacts (intrusions) with blue nodes indicating males, yellow nodes indicating females, and gray nodes indicating non-focal squirrels (i.e., those without microbiota samples); edge thickness is intrusion-based social association index, corrected for lifespan overlap. **(B)** Social association strength predicts gut microbial similarity, controlling for spatial distance between pairs. Forest plot shows posterior means (points) and their 95% credible intervals from Bayesian regression model using gut microbiome similarity (Jaccard Index) between pairs as the response variable. Credible intervals (CIs) that do not overlap zero indicate factors that significantly predicted gut microbial similarity. The effect of read depth difference removed from the plot for better visualization of CIs but retained in the model (see Table S13). (**C)** Slope of the effect of pairwise social association on gut microbial similarity (conditional effects with 95% CIs).

## DISCUSSION

Social behaviors that facilitate the transmission of microbiota across hosts may confer individual and/or population resilience by enhancing host microbial diversity and flexibility, particularly in changing environments [13, 20, 38]. In highly territorial species in which individuals maintain social isolation through spatial and behavioral mechanisms, agonistic interactions may facilitate social microbial transmission, or the social transfer of microbiota may be absent entirely. Here, we find that elevated population densities and territorial intrusions diversify and homogenize gut microbial communities in territorial red squirrels, providing the first evidence for social microbial transmission in a non-social species.

Microbial evolution within the host gastrointestinal tract can be rapid, generating genomically distinct variants of microbial taxa within a host’s lifetime [39], increasing microbial diversity [40]. When population densities are high, rates of incidental physical contact among conspecifics can increase, and opportunities for microbial transmission are likely to be greater [34]. Indeed, when squirrel densities increased, individual gut microbial communities became taxonomically richer and phylogenetically more diverse, echoing prior work in highly social pikas *(Ochotona curzoniae)* suggestive of social microbial transmission [41]. This increase in diversity is likely independent of dietary diversification: squirrel densities peak following a spruce mast when dietary heterogeneity shrinks as squirrels primarily consume their new larder hoard of spruce cones [28, 29]. It is also unlikely to result from physiological responses to conspecific density, as we have previously shown that the physiological stress response (indicated by elevated fecal glucocorticoid metabolites) to increased density decreases microbial diversity [42]. Still, if the diversification of individual gut microbiota was unrelated to microbial transmission, elevated densities should exacerbate individual differences in microbial composition [34]. Instead, microbial similarity among squirrels was highest during periods of high density, and decreased as densities shrunk, suggesting microbial homogenization through host-to-host transmission [34].

Most of the microbial genera that were differentially enriched with increasing squirrel densities were obligately anaerobic and non-sporulating taxa unlikely to survive long outside of the anoxic gut environment. For example, at higher squirrel densities, the gut microbiota was enriched in *Gastranaerophilales,* the only order of cyanobacteria-related Melainabacteria lacking genes for aerobic respiration [43, 44], commonly found in the gastrointestinal tracts of humans and other animals [45]. Other enriched obligate anaerobes included *Herbinix spp.,* non-sporulating cellulose degraders whose functional roles in the host microbiome are not yet well understood [46], and the butyrate-producing genera *Frisingicoccus,* involved in host metabolism and ketogenesis [47]. The social transfer of these and other obligately anaerobic bacteria is thought to require intimate host-to-host interactions [35, 40], but physical contact between red squirrels is rarely observed [48]. Transmission across individuals may instead occur indirectly through fecal-oral routes of transmission like defecation and coprophagy, which is common in rodents [38, 49]. In line with this interpretation, the largest effect of increasing density was an increase in *Sanguibacter*, a genus of facultative anaerobes with some aerotolerance and thus capability for survival between hosts [50]. Together, shifts in obligate anaerobes may indicate some intimate host-to-host transmission during periods of high density, but in such a non-social species, the most successful transmission may occur within taxa with at least some capacity for survival in an environmental in-between. Nonetheless, the overall lack of aerobic and spore-forming taxa in the red squirrel gut microbiota may reflect limited evolutionary selection for bacterial traits that enhance transmission in a species with infrequent social interactions [19].

The intimate symbiosis between hosts and their microbiota generates microbial communities whose primary functions fluctuate with the physiological demands of their host [51]. At lower squirrel densities, gut microbiota expressed greater capacity for functions related to fundamental host physiological processes including metabolism, immunity, digestion, and endocrine function. At higher squirrel densities, gut microbiota shifted toward a greater capacity for the processing of environmental information. Pathways related to membrane transport and signal transduction became more abundant, as did pathways related to environmental adaptation, including thermogenesis and circadian rhythm entrainment. This shift from “inward” (e.g., involved in major organ and physiological systems) to “outward” (e.g., involved in environmental sensing) functions may reflect host prioritization of microbiota with the capacity to detect and integrate environmental cues when intraspecific competition is high. It may also reflect the displacement of core taxa (e.g., *Prevotella spp.,* [52]) central to the regulation of intrinsic host physiological systems as a result of the stochastic incorporation of socially-acquired microbiota [14].

Applications of island biogeography theory to microbial ecology typically focus on hosts as microbial islands [53], but non-overlapping, individual territories can be similarly viewed as islands. Principles of species-area relationships thus predict that the microbial diversity within a given territory should increase with increasing territory size [54]. If the effect of territory size on gut microbiota was driven by environmental transmission, we would expect individuals defending larger territories to house a more diverse microbiota. Instead, squirrels defending smaller territories exhibited greater microbial diversity. Moreover, more intrusions were associated with greater microbial diversity, but only among the owners of large territories. Territory size and the frequency of intrusions are positively correlated in red squirrels [32], suggesting a potential threshold effect by which detectable signatures of intrusion-mediated changes to microbial communities depend on a higher baseline of opportunities for social microbial transmission.

In highly social animals, strong social bonds that promote affiliative interactions like grooming can enrich microbial communities [55]. Our data suggest that territorial intrusions have a similar diversifying effect, though perhaps only for individuals defending territories highly susceptible to intrusions. Nonetheless, the diversifying effect of higher squirrel densities and more frequent territorial intrusions may confer substantial benefits for both individual fitness and population persistence. The maintenance of microbial diversity is crucial to pathogen defense as it increases the competitive nutritional environment in the gut, blocking invading pathogens from successfully colonizing the host gastrointestinal tract [56]. At the population level, a diverse pool of gut microbiota can buffer against random microbial extinction events associated with drift [13]. If extinctions do occur, microbial function can nonetheless be preserved through functional redundancy conferred by a diverse set of microbes [57, 58].

In social and semi-social species, the stronger the social relationship between a pair of individuals, the more similar their gut microbiota [17, 23]. Here, we additionally uncover a parallel effect in an asocial mammal when measuring social association strength based on agonistic interactions between two squirrels. By generating a measure of social association among pairs of squirrels based on the relative number of pairwise intrusions between them, we found that pairs with more intrusions had more similar microbiota, controlling for spatial proximity (distance between territory centroids). This finding suggests that pairs of squirrels with territorial intrusions between them exhibit some degree of social microbial transmission, though the route of transfer (direct via physical contact vs. indirect via fecal shedding) remains unknown. Given the risks of pilferage and energetic stress associated with territorial intrusions, we hypothesize that the homogenization of microbial communities via intrusions may be a cryptic benefit that could reduce the cost of intruding. Intrusions can weaken the stability of territorial systems by making resources less valuable and defense more costly [59], but if they facilitate the spread of beneficial bacteria across otherwise non-interacting hosts, they may confer benefits to both the owner and intruder.

In sum, we provide evidence for territorial behavior as a means of social microbial transmission in a non-social mammal. Our findings suggest that the underlying biological mechanisms facilitating the incorporation of socially-transmitted microbiota may be maintained even in species where physical contact is rare. While the exact nature of such mechanisms remains unclear, our findings suggest that the social transfer of commensal microbiota is not unique to social and semi-social species, and that agonistic behaviors may also facilitate microbial transfer. As such, our results suggest that the host-to-host transfer of microbial symbionts may be an inconspicuous benefit of territorial behaviors and may provide insight into the resilience of microbial transmission pathways during periods of social isolation and disconnect in more social species.

## ACKNOWLEDGMENTS

We thank Agnes MacDonald and her family for long-term access to her trapline, and the Champagne and Aishihik First Nations for allowing us to conduct our work within their traditional territory. We especially thank all of the field technicians that contributed to data collection, and Pippin Ke Lind and John Paul Williams-Soriano for assistance with wet lab work. We also wish to thank Animal Microbiome Research Group network for bringing some of the authors together. Sequencing was performed by the University of Michigan Microbiome Core. Bioinformatic analyses used the High-Performance Computing (HPC) resources supported by the University of Arizona TRIF, UITS, and Research, Innovation, and Impact (RII) and maintained by the University of Arizona Research Technologies department. This work was supported by the University of Arizona, the University of Michigan, the National Science Foundation (DEB-2010726 to LP, DEB-0515849 to AGM, IOS-1749627 to BD, and DEB-2338394 to AGM and BD) and the Natural Sciences and Engineering Research Council of Canada to (SB, AGM, JEL, QW). This is KRSP paper #XXX.

## DATA AVAILABILITY STATEMENT

All data and code associated with this manuscript will be made publicly and freely accessible at the following FigShare repository at the time of publication: https://figshare.com/s/a79d2e1603fe55ac91ce. The code for social network analysis and Bayesian dyadic models can be found from AR’s github page: https://github.com/nuorenarra. Raw sequence data has been submitted to NCBI Sequence Read Archive.

