## Supplemental Materials for "Territorial behavior as a route of social microbial transmission in an asocial mammal"

### Table S1. Temporal and methodological factors explaining variation in gut microbial community composition. Results from a marginal PERMANOVA on the entire dataset (52 individuals, 562 samples) with Jaccard Index as the response variable. N = 999 permutations.

|  | **Df** | **R2** | **F** | ***p*** |
| --- | --- | --- | --- | --- |
| **Individual ID** | **51** | **0.195** | **2.819** | **0.001** |
| **Season** | **3** | **0.030** | **7.399** | **0.001** |
| **Year** | **8** | **0.041** | **3.722** | **0.001** |
| Sequencing run | 1 | 0.001 | 1.004 | 0.448 |
| **Sample read depth** | **1** | **0.013** | **9.438** | **0.001** |
| Residual | 497 |  |  |  |
| Total | 561 |  |  |  |

### Table S2. Intrinsic and spatial host factors explaining variation in gut microbial community composition, controlling for methodological and temporal factors. Results from a marginal PERMANOVA on the entire dataset (52 individuals, 562 samples) with Jaccard Index as the response variable. N = 999 permutations, individual ID set as blocking factor using permute.

|  | **Df** | **R2** | **F** | ***p*** |
| --- | --- | --- | --- | --- |
| Age | 1 | 0.004 | 2.425 | 0.871 |
| Sex | 1 | 0.003 | 1.695 | 0.388 |
| **Study area** | **1** | **0.006** | **3.919** | **0.001** |
| **Season** | **3** | **0.025** | **5.048** | **0.001** |
| **Year** | **8** | **0.042** | **3.164** | **0.001** |
| **Sample read depth** | **1** | **0.009** | **5.410** | **0.001** |
| Residual | 546 | 0.901 |  |  |
| Total | 561 | 1 |  |  |

### Table S3. Gut microbial individuality fluctuates with fluctuating squirrel density. Results from a generalized linear mixed-effects model (family = “beta”) testing the effects of squirrel density (high, medium, low, based on a three-way split of total study area density) on distance to group centroid in gut microbiota compositional space (Jaccard similarity). Model controls for confounding effects of study area (KL or SU), sampling season, and sample read depth.

|  | **Estimate** | **Std. Error** | **z** | ***p*** |
| --- | --- | --- | --- | --- |
| **Density (medium)** | **-0.046** | **0.014** | **-3.336** | **0.000** |
| **Density (high)** | **-0.067** | **0.020** | **-3.324** | **0.000** |
| Study area (SU) | 0.006 | 0.018 | 0.319 | 0.750 |
| Season (fall) | -0.026 | 0.029 | -0.919 | 0.358 |
| **Season (late spring)** | **0.024** | **0.009** | **2.538** | **0.011** |
| **Season (summer)** | **0.033** | **0.013** | **2.792** | **0.005** |
| **Read depth** | **-0.057** | **0.004** | **-14.505** | **0.000** |

**Table S4.** **Neighborhood squirrel densities predict individual gut microbial alpha-diversity to different degrees.** Results from LMMs/GLMMs testing the effect of neighborhood density (within 50 m of an individual’s territory) on three measures of gut microbial alpha-diversity.

| **A** | **ASV richness** | | | | | |
| --- | --- | --- | --- | --- | --- | --- |
|  |  | **Estimate** | **Std error** | **df** | **t** | ***p*** |
|  | **Neighborhood density (within 50 m)** | **4.222** | **1.962** | **459.805** | **2.152** | **0.032** |
|  | Mast (no) | -22.580 | 10.711 | 8.448 | -2.108 | 0.066 |
|  | Sex (male) | -2.013 | 7.306 | 54.796 | -0.275 | 0.784 |
|  | Season (fall) | 14.297 | 11.417 | 543.078 | 1.252 | 0.211 |
|  | **Season (late spring)** | **-27.024** | **3.958** | **542.072** | **-6.828** | **0.000** |
|  | Season (summer) | -5.732 | 4.808 | 534.508 | -1.192 | 0.234 |
|  | Sequencing run | 5.607 | 6.616 | 530.181 | 0.847 | 0.397 |
|  | Study area (SU) | -3.179 | 5.743 | 48.145 | -0.554 | 0.583 |
| **B** | **Shannon Index (abundance weighted)** | | | | | |
|  | **Neighborhood density (within 50 m)** | **52.962** | **15.656** | **540.052** | **3.383** | **0.001** |
|  | Mast (no) | -193.727 | 115.54 | 7.591 | -1.677 | 0.134 |
|  | Sex (male) | 18.285 | 60.846 | 57.109 | 0.301 | 0.765 |
|  | Season (fall) | -17.69 | 89.907 | 539.695 | -0.197 | 0.844 |
|  | **Season (late spring)** | **-298.133** | **31.231** | **544.234** | **-9.546** | **0.000** |
|  | **Season (summer)** | **-140.714** | **37.96** | **542.545** | **-3.707** | **0.000** |
|  | Sequencing run | 50.323 | 51.915 | 525.569 | 0.969 | 0.333 |
|  | Study area (SU) | -58.141 | 47.635 | 48.193 | -1.221 | 0.228 |
| **C** | **Faith's Phylogenetic diversity** | | | | | |
|  | **Neighborhood density (within 50 m)** | **0.159** | **0.080** | **528.731** | **1.992** | **0.047** |
|  | Mast (no) | -0.888 | 0.538 | 7.995 | -1.651 | 0.137 |
|  | Sex (male) | -0.089 | 0.321 | 58.299 | -0.276 | 0.784 |
|  | Season (fall) | -0.223 | 0.459 | 539.286 | -0.485 | 0.628 |
|  | **Season (late spring)** | **-0.901** | **0.160** | **542.480** | **-5.646** | **0.000** |
|  | Season (summer) | -0.311 | 0.194 | 539.879 | -1.601 | 0.110 |
|  | Sequencing run | -0.133 | 0.265 | 525.632 | -0.503 | 0.615 |
|  | Study area (SU) | -0.038 | 0.253 | 50.132 | -0.150 | 0.882 |

### Table S5. Results from generalized/linear-mixed effects models testing the effect of neighborhood density (within 100 m and 150 m of an individual’s territory) on three measures of gut microbial alpha-diversity.

| **A** | **ASV richness** | | | | | |
| --- | --- | --- | --- | --- | --- | --- |
|  |  | **Estimate** | **Std error** | **df** | **t** | ***p*** |
|  | Neighborhood density (within 100 m) | 3.464 | 1.993 | 405.200 | 1.738 | 0.083 |
|  | Mast (no) | -22.180 | 10.540 | 8.584 | -2.106 | 0.066 |
|  | Sex (male) | -2.141 | 7.340 | 53.960 | -0.292 | 0.772 |
|  | Season (fall) | 14.210 | 11.440 | 542.200 | 1.242 | 0.215 |
|  | **Season (late spring)** | **-26.770** | **4.032** | **539.800** | **-6.639** | **0.000** |
|  | Season (summer) | -5.444 | 4.813 | 528.500 | -1.131 | 0.259 |
|  | Sequencing run | 5.478 | 6.627 | 530.400 | 0.827 | 0.409 |
|  | Study area (SU) | -3.030 | 5.837 | 0.000 | -0.519 | 1.000 |
|  | Neighborhood density (within 150 m) | 3.771 | 2.036 | 346.602 | 1.852 | 0.065 |
|  | Mast (no) | -22.445 | 10.335 | 8.639 | -2.172 | 0.059 |
|  | Sex (male) | -2.237 | 7.325 | 53.445 | -0.305 | 0.761 |
|  | Season (fall) | 14.120 | 11.438 | 542.194 | 1.234 | 0.218 |
|  | **Season (late spring)** | **-27.176** | **4.087** | **538.027** | **-6.649** | **0.000** |
|  | Season (summer) | -5.599 | 4.814 | 524.638 | -1.163 | 0.245 |
|  | Sequencing run | 5.665 | 6.623 | 530.171 | 0.855 | 0.393 |
|  | Study area (SU) | -3.183 | 5.766 | 47.487 | -0.552 | 0.584 |
| **B** | **Shannon Index (abundance weighted)** | | | | | |
|  | **Neighborhood density (within 100 m)** | **50.971** | **15.944** | **524.009** | **3.197** | **0.001** |
|  | Mast (no) | -189.998 | 112.711 | 7.600 | -1.686 | 0.132 |
|  | Sex (male) | 15.927 | 61.460 | 56.688 | 0.259 | 0.796 |
|  | Season (fall) | -20.968 | 90.016 | 538.751 | -0.233 | 0.816 |
|  | **Season (late spring)** | **-300.693** | **31.782** | **540.871** | **-9.461** | **0.000** |
|  | **Season (summer)** | **-140.050** | **37.995** | **539.688** | **-3.686** | **0.000** |
|  | Sequencing run | 47.593 | 51.987 | 525.988 | 0.915 | 0.360 |
|  | Study area (SU) | -55.276 | 48.206 | 48.249 | -1.147 | 0.257 |
|  | **Neighborhood density (within 150 m)** | **46.311** | **16.400** | **507.058** | **2.824** | **0.005** |
|  | Mast (no) | -193.002 | 112.431 | 7.612 | -1.717 | 0.126 |
|  | Sex (male) | 14.970 | 61.279 | 56.506 | 0.244 | 0.808 |
|  | Season (fall) | -19.553 | 90.226 | 538.970 | -0.217 | 0.829 |
|  | **Season (late spring)** | **-299.787** | **32.325** | **541.656** | **-9.274** | **0.000** |
|  | **Season (summer)** | **-138.806** | **38.119** | **539.483** | **-3.641** | **0.000** |
|  | Sequencing run | 51.273 | 52.075 | 525.859 | 0.985 | 0.325 |
|  | Study area (SU) | -58.459 | 47.990 | 47.917 | -1.218 | 0.229 |
| **C** | **Faith's Phylogenetic Diversity** | | | | | |
|  | **Neighborhood density (within 100 m)** | **0.179** | **0.081** | **502.437** | **2.210** | **0.028** |
|  | Mast (no) | -0.878 | 0.526 | 8.018 | -1.669 | 0.134 |
|  | Sex (male) | -0.097 | 0.323 | 57.677 | -0.300 | 0.765 |
|  | Season (fall) | -0.239 | 0.459 | 538.408 | -0.521 | 0.602 |
|  | **Season (late spring)** | **-0.927** | **0.162** | **539.328** | **-5.719** | **0.000** |
|  | Season (summer) | -0.317 | 0.194 | 536.635 | -1.638 | 0.102 |
|  | Sequencing run | -0.145 | 0.265 | 526.026 | -0.547 | 0.584 |
|  | Study area (SU) | -0.024 | 0.254 | 49.941 | -0.095 | 0.925 |
|  | **Neighborhood density (within 150 m)** | **0.170** | **0.083** | **477.409** | **2.040** | **0.042** |
|  | Mast (no) | -0.889 | 0.524 | 8.013 | -1.697 | 0.128 |
|  | Sex (male) | -0.101 | 0.323 | 57.503 | -0.313 | 0.755 |
|  | Season (fall) | -0.236 | 0.459 | 538.422 | -0.514 | 0.607 |
|  | **Season (late spring)** | **-0.929** | **0.165** | **539.503** | **-5.643** | **0.000** |
|  | Season (summer) | -0.315 | 0.194 | 535.962 | -1.623 | 0.105 |
|  | Sequencing run | -0.133 | 0.265 | 525.690 | -0.502 | 0.616 |
|  | Study area (SU) | -0.035 | 0.254 | 49.686 | -0.136 | 0.892 |

### Table S6. At the level of an individual’s entire study area, squirrel density did not predict gut microbial alpha-diversity. Results from generalized/linear mixed-effects models testing the effect of squirrel population densities within an individual’s entire study area (~40 ha) on three measures of gut microbial alpha-diversity.

| **A** | **ASV richness** | | | | | |
| --- | --- | --- | --- | --- | --- | --- |
|  | **Variable** | **Estimate** | **Std error** | **df** | **t** | **P** |
|  | Squirrel density (study area) | -1.369 | 4.011 | 6.873 | -0.341 | 0.743 |
|  | Mast (no) | -20.497 | 12.968 | 6.336 | -1.581 | 0.162 |
|  | Sex (male) | -1.518 | 7.485 | 58.613 | -0.203 | 0.840 |
|  | Season (fall) | 15.154 | 11.457 | 542.474 | 1.323 | 0.187 |
|  | Season (late spring) | -24.286 | 3.748 | 536.304 | -6.481 | 0.000 |
|  | Season (summer) | -4.105 | 4.784 | 539.142 | -0.858 | 0.391 |
|  | Sequencing run | 0.018 | 0.020 | 527.920 | 0.933 | 0.351 |
|  | Study area (SU) | -4.013 | 5.947 | 52.313 | -0.675 | 0.503 |
| **B** | **Shannon Index (abundance weighted)** | | | | | |
|  | Squirrel density (study area) | 5.405 | 43.747 | 6.662 | 0.124 | 0.905 |
|  | Mast (no) | -197.758 | 142.753 | 6.219 | -1.385 | 0.214 |
|  | Sex (male) | 22.117 | 61.705 | 58.421 | 0.358 | 0.721 |
|  | Season (fall) | -4.733 | 90.834 | 537.618 | -0.052 | 0.958 |
|  | Season (late spring) | -263.227 | 29.745 | 540.849 | -8.849 | 0.000 |
|  | Season (summer) | -119.578 | 37.967 | 543.490 | -3.150 | 0.002 |
|  | Sequencing run | 0.167 | 0.156 | 524.493 | 1.073 | 0.284 |
|  | Study area (SU) | -61.694 | 50.432 | 54.250 | -1.223 | 0.227 |
| **C** | **Faith's Phylogenetic diversity** | | | | | |
|  | Squirrel density (study area) | 0.122 | 0.190 | 6.942 | 0.643 | 0.541 |
|  | Mast (no) | -1.029 | 0.614 | 6.421 | -1.675 | 0.142 |
|  | Sex (male) | -0.107 | 0.325 | 59.880 | -0.327 | 0.744 |
|  | Season (fall) | -0.178 | 0.461 | 538.500 | -0.387 | 0.699 |
|  | Season (late spring) | -0.799 | 0.151 | 537.500 | -5.294 | 0.000 |
|  | Season (summer) | -0.244 | 0.193 | 540.100 | -1.265 | 0.206 |
|  | Sequencing run | 0.000 | 0.001 | 524.200 | -0.466 | 0.641 |
|  | Study area (SU) | -0.013 | 0.262 | 48.840 | -0.051 | 0.959 |

### Table S7. Ten microbial genera significantly vary in relative abundance with squirrel densities. Results depict significant (*p_FDR_* < 0.05) terms from zero-inflated negative binomial mixed-effects models testing the effects of neighborhood density (within 50 m) on the relative abundance of bacterial genera (modeled as raw counts), controlling for methodological and ecological covariates. Sample read depth included as log-transformed model offset and individual identity and year as random effects.

| **Microbial genus** | **Estimate** | **p** | ***p* (FDR-adjusted)** |
| --- | --- | --- | --- |
| Gastranaerophilales | 0.100 | 0.003 | 0.035 |
| Prevotellaceae_UCG-001 | -0.135 | 0.000 | 0.011 |
| Oscillospiraceae_UCG-005 | 0.084 | 0.003 | 0.035 |
| Prevotellaceae_Ga6A1_group | -0.153 | 0.004 | 0.040 |
| Oscillospiraceae_NK4A214_group | 0.089 | 0.006 | 0.047 |
| Erysipelatoclostridium | 0.132 | 0.000 | 0.004 |
| Sanguibacter | 0.527 | 0.006 | 0.047 |
| Frisingicoccus | 0.167 | 0.005 | 0.047 |
| Eubacterium_nodatum_group | 0.276 | 0.000 | 0.000 |
| Herbinix | 0.342 | 0.001 | 0.026 |

### Table S8. Predicted functional pathways at Level II of the BRITE map that are differentially enriched in the gut microbiota as a function of neighborhood density (within 50 m). Results depict significant (*p_FDR_* < 0.05) terms from a series of linear mixed-effects models testing whether density predicts differences in the relative abundance of functional pathways, controlling for mast year (yes/no), sex, sampling season, sequencing run, and study area as fixed factors, with year and individual ID as random effects.

| **Functional pathway (Level 2)** | **Estimate** | ***p*** | ***p* (FDR-adjusted)** |
| --- | --- | --- | --- |
| Biosynthesis of Other Secondary Metabolites | -0.006 | 0.001 | 0.007 |
| Digestive System | -0.003 | 0.000 | 0.003 |
| Endocrine System | -0.003 | 0.000 | 0.006 |
| Energy Metabolism | -0.024 | 0.001 | 0.008 |
| Environmental Adaptation | 0.001 | 0.013 | 0.046 |
| Glycan Biosynthesis and Metabolism | -0.032 | 0.003 | 0.011 |
| Immune System Diseases | -0.001 | 0.002 | 0.008 |
| Membrane Transport | 0.078 | 0.014 | 0.046 |
| Metabolism | -0.005 | 0.015 | 0.046 |
| Metabolism of Other Amino Acids | -0.005 | 0.008 | 0.032 |
| Signal Transduction | 0.015 | 0.002 | 0.008 |
| Signaling Molecules and Interaction | -0.002 | 0.002 | 0.008 |
| Transport and Catabolism | -0.006 | 0.000 | 0.003 |

### Table S9. Territorial spaces fluctuate in size as a function of intrinsic and extrinsic factors. Results from linear mixed-effects model investigating predictors of variation in the size of an individual’s territorial space. Model included individual ID and year as random effects.

|  | **Estimate** | **Std error** | **df** | **t** | ***p*** |
| --- | --- | --- | --- | --- | --- |
| **Neighborhood density (within 50 m)** | **-0.022** | **0.011** | **517.080** | **-2.014** | **0.045** |
| Mast (no) | -0.237 | 0.128 | 5.834 | -1.847 | 0.116 |
| **Sex (male)** | **0.266** | **0.114** | **50.383** | **2.328** | **0.024** |
| Season (fall) | 0.037 | 0.060 | 509.888 | 0.617 | 0.537 |
| **Season (late spring)** | **0.059** | **0.021** | **508.762** | **2.830** | **0.005** |
| Season (summer) | 0.006 | 0.025 | 508.803 | 0.228 | 0.820 |
| Study area (SU) | -0.045 | 0.094 | 48.598 | -0.479 | 0.634 |

### Table S10. The rate of territorial intrusions is greater among squirrels with larger territorial spaces. Results from linear mixed-effects model testing the relationship between the area of an individual’s territorial space (ha) and frequency of intrusions (dependent variable), controlling for potential confounding variables. Model included individual ID and year as random effects.

|  | **Estimate** | **Std error** | **df** | **t** | **p** |
| --- | --- | --- | --- | --- | --- |
| **Area (ha)** | **0.063** | **0.007** | **548.100** | **9.583** | **0.000** |
| Neighborhood density (within 50 m) | -0.001 | 0.005 | 510.800 | -0.168 | 0.866 |
| Mast (no) | -0.167 | 0.105 | 4.552 | -1.590 | 0.178 |
| Sex (male) | -0.050 | 0.050 | 52.900 | -0.989 | 0.327 |
| **Season (fall)** | **-0.054** | **0.027** | **509.100** | **-2.022** | **0.044** |
| Season (late spring) | 0.010 | 0.009 | 503.400 | 1.035 | 0.301 |
| Season (summer) | -0.004 | 0.011 | 502.400 | -0.340 | 0.734 |
| **Study area (SU)** | **-0.125** | **0.041** | **50.030** | **-3.059** | **0.004** |

### Table S11. Relationship between territorial area, territorial intrusions, and gut microbial alpha-diversity. All models controlled for differences in sample read depth (log-transformed and included as a model offset), and random effects of year and individual identity. Significant (*p* < 0.05) terms shown in bold.

| A | ASV richness | | | | | |
| --- | --- | --- | --- | --- | --- | --- |
|  |  | Estimate | Std error | df | t | P |
|  | Intrusion frequency | 2.759 | 2.696 | 145.965 | 1.024 | 0.308 |
|  | Area (ha) | -11.698 | 3.030 | 186.624 | -3.861 | 0.000 |
|  | Neighborhood density (within 50 m) | 4.092 | 1.940 | 439.496 | 2.109 | 0.035 |
|  | Mast (no) | -22.878 | 10.880 | 10.444 | -2.103 | 0.061 |
|  | Sex (male) | 0.044 | 7.267 | 56.292 | 0.006 | 0.995 |
|  | Season (fall) | 16.688 | 11.351 | 542.220 | 1.470 | 0.142 |
|  | Season (late spring) | -25.963 | 3.935 | 531.945 | -6.598 | 0.000 |
|  | Season (summer) | -3.653 | 4.797 | 535.012 | -0.762 | 0.447 |
|  | Sequencing run | 4.473 | 6.564 | 530.027 | 0.681 | 0.496 |
|  | Study area (SU) | -1.545 | 5.658 | 47.632 | -0.273 | 0.786 |
|  | Intrusion frequency x area (ha) | 6.276 | 2.026 | 311.859 | 3.097 | 0.002 |
| B | Shannon Index (abundance weighted) | | | | | |
|  | Intrusion frequency | 23.687 | 22.007 | 159.458 | 1.076 | 0.283 |
|  | Area (ha) | -87.960 | 24.622 | 205.276 | -3.572 | 0.000 |
|  | Neighborhood density (within 50 m) | 51.368 | 15.528 | 525.920 | 3.308 | 0.001 |
|  | Mast (no) | -198.549 | 112.153 | 8.479 | -1.770 | 0.113 |
|  | Sex (male) | 44.173 | 60.401 | 61.385 | 0.731 | 0.467 |
|  | Season (fall) | -1.846 | 89.624 | 539.677 | -0.021 | 0.984 |
|  | Season (late spring) | -288.822 | 31.195 | 540.562 | -9.259 | 0.000 |
|  | Season (summer) | -128.788 | 37.930 | 538.598 | -3.395 | 0.001 |
|  | Sequencing run | 43.194 | 51.686 | 526.061 | 0.836 | 0.404 |
|  | Study area (SU) | -42.599 | 46.678 | 48.051 | -0.913 | 0.366 |
|  | Intrusion frequency x area (ha) | 41.071 | 16.267 | 316.850 | 2.525 | 0.012 |
| C | Faith's Phylogenetic Diversity | | | | | |
|  | Intrusion frequency | 0.129 | 0.114 | 183.625 | 1.123 | 0.263 |
|  | Area (ha) | -0.449 | 0.128 | 222.575 | -3.515 | 0.001 |
|  | Neighborhood density (within 50 m) | 0.161 | 0.079 | 497.905 | 2.040 | 0.042 |
|  | Mast (no) | -0.840 | 0.498 | 10.027 | -1.687 | 0.122 |
|  | Sex (male) | -0.085 | 0.321 | 61.441 | -0.264 | 0.792 |
|  | Season (fall) | -0.104 | 0.456 | 538.590 | -0.227 | 0.820 |
|  | Season (late spring) | -0.865 | 0.159 | 535.103 | -5.451 | 0.000 |
|  | Season (summer) | -0.231 | 0.193 | 534.612 | -1.200 | 0.231 |
|  | Sequencing run | -0.183 | 0.263 | 525.641 | -0.694 | 0.488 |
|  | Study area (SU) | 0.048 | 0.251 | 50.616 | 0.189 | 0.850 |
|  | Intrusion frequency x area (ha) | 0.316 | 0.084 | 337.111 | 3.761 | 0.000 |

### Table S12. Predictors of gut microbial similarity across all pairs of squirrels. Results from *brms* model testing the effects of potential covariates on Jaccard similarity across squirrel pairs (N = 1,326 pairs). Significant (where 95% credible intervals do not overlap zero) terms shown in bold.

|  | **Estimate** | **Est.error** | **l-95% CI** | **u-95% CI** | **Rhat** | **Bulk_ESS** | **Tail_ESS** |
| --- | --- | --- | --- | --- | --- | --- | --- |
| Same sex | 0 | 0 | 0 | 0 | 1 | 6852 | 9879 |
| **Same study area** | **0.02** | **0** | **0.01** | **0.02** | **1** | **11012** | **15423** |
| **Same year** | **0.1** | **0** | **0.1** | **0.1** | **1** | **7063** | **11085** |
| **Same season** | **0.14** | **0** | **0.14** | **0.14** | **1** | **10496** | **14551** |
| **Age difference** | **-0.01** | **0** | **-0.02** | **0** | **1** | **5478** | **9448** |
| **Read depth difference** | **-0.86** | **0.01** | **-0.88** | **-0.85** | **1** | **4599** | **8134** |

### Table S13. Social association strength predicts gut microbiota similarity among pairs of squirrels inhabiting the same study area and sampled in the same year. Results from *brms* model testing the effects of social association (intrusions) on Jaccard similarity across squirrel pairs (N = 436 pairs). Significant (where 95% credible intervals do not overlap zero) terms shown in bold.

|  | **Estimate** | **Est. error** | **l-95% CI** | **u-95% CI** | **Rhat** | **Bulk_ESS** | **Tail_ESS** |
| --- | --- | --- | --- | --- | --- | --- | --- |
| **Same sex** | **0** | **0** | **0** | **0.01** | **1** | **4722** | **2558** |
| **Same season** | **0.18** | **0** | **0.18** | **0.19** | **1** | **4564** | **2969** |
| Age difference | 0 | 0.01 | -0.02 | 0.01 | 1 | 4835 | 2867 |
| **Read depth difference** | **-0.79** | **0.01** | **-0.81** | **-0.77** | **1** | **4563** | **2631** |
| **Spatial distance** | **-0.1** | **0.01** | **-0.12** | **-0.08** | **1** | **4385** | **3370** |
| **Social association** | **0.22** | **0.04** | **0.14** | **0.29** | **1** | **4624** | **2911** |

### Figure S1. Homogeneity of dispersion in gut microbial communities varies as a function of population density. Points (samples) and group centroids (medians) in a principal-coordinates derived euclidean space based on Jaccard similarity, colored by squirrel density within the entire ~40 ha study area (low density = black, medium density = purple, high density = pink; based on an even three-way split). Confidence ellipses reflect 1 standard deviation from the median.

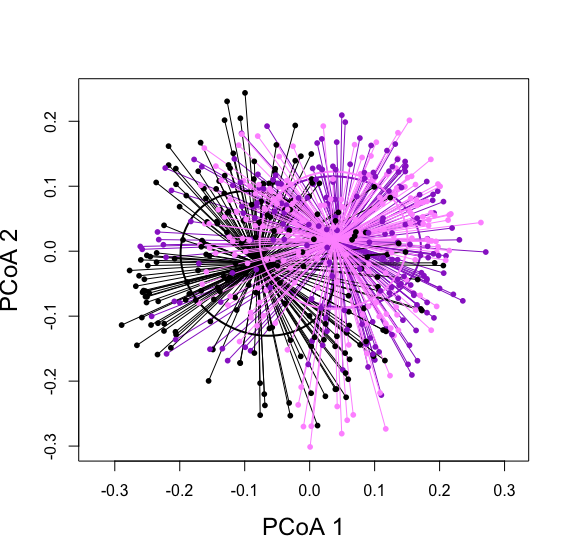

### Figure S2. Appreciable variation in squirrel densities at coarse and fine scales on both study areas across the sampling period. Dotted lines reflect white spruce mast (food pulse) years, in which spring population densities are typically at their lowest and sharply increase the following year as a result of a superabundance of food.

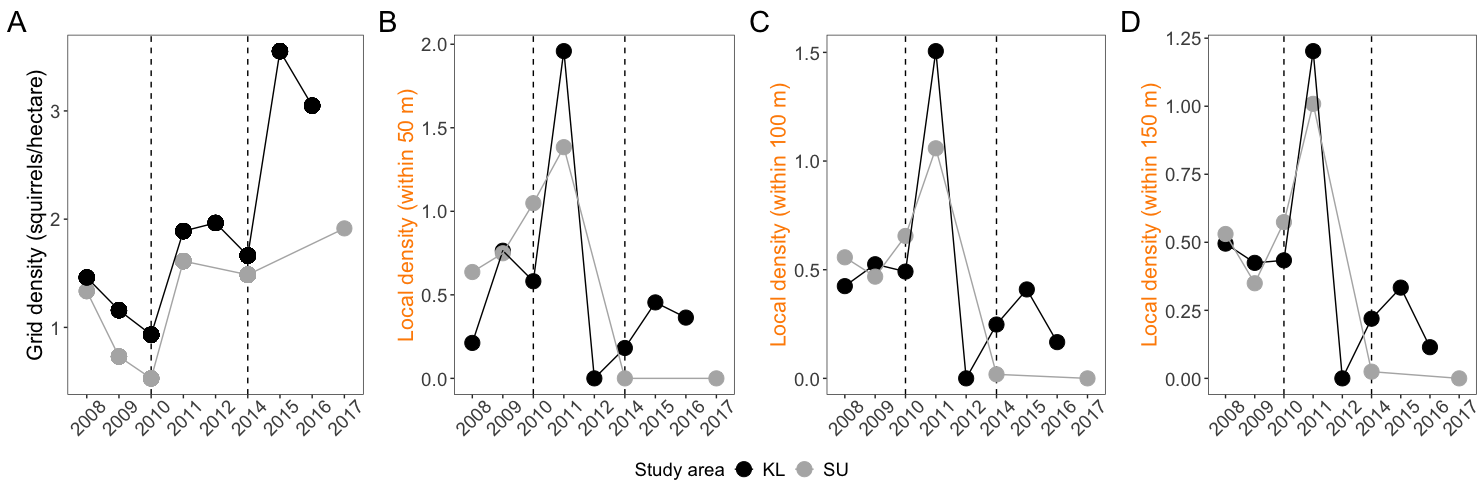

### Figure S3. Correlations among different measures of squirrel population densities across years of the study period.

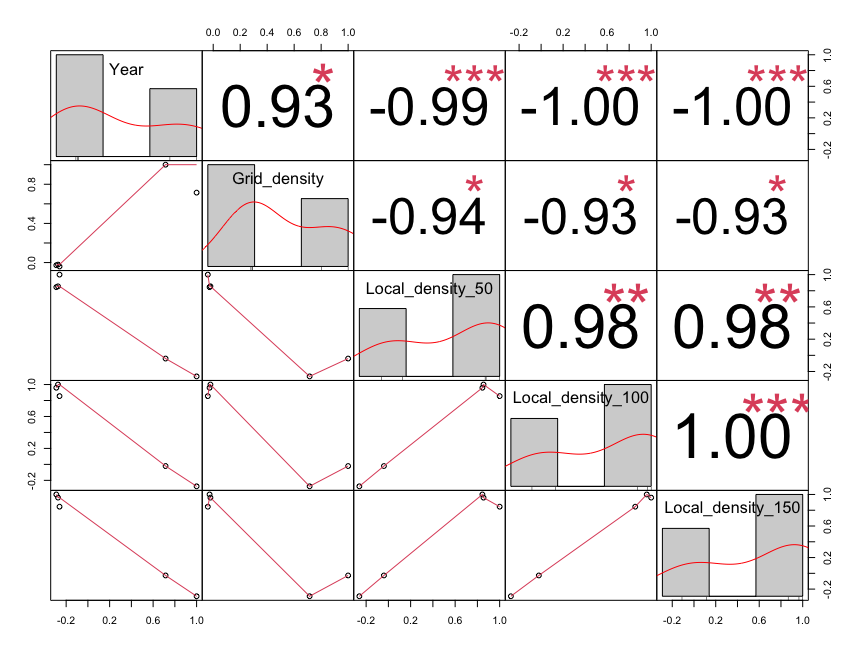

**Figure S4. Survey of predicted functional microbial pathways present within the red squirrel gut microbiota.** The majority of functional pathways identified using the PICRUSt2 pipeline in the red squirrel gut microbiota were related to membrane transport (median relative abundance = 10.4%, range = 8.2 - 12.7%), replication and repair (median relative abundance = 10.3%, range = 9.6 - 11.0%), and carbohydrate (median relative abundance = 9.7%, range 9.3 - 10.8%) and amino acid metabolism (median relative abundance = 9.7%, range = 9.3-10.4%). Pathways depicted below identified at Level II of the BRITE map across all samples in the dataset.

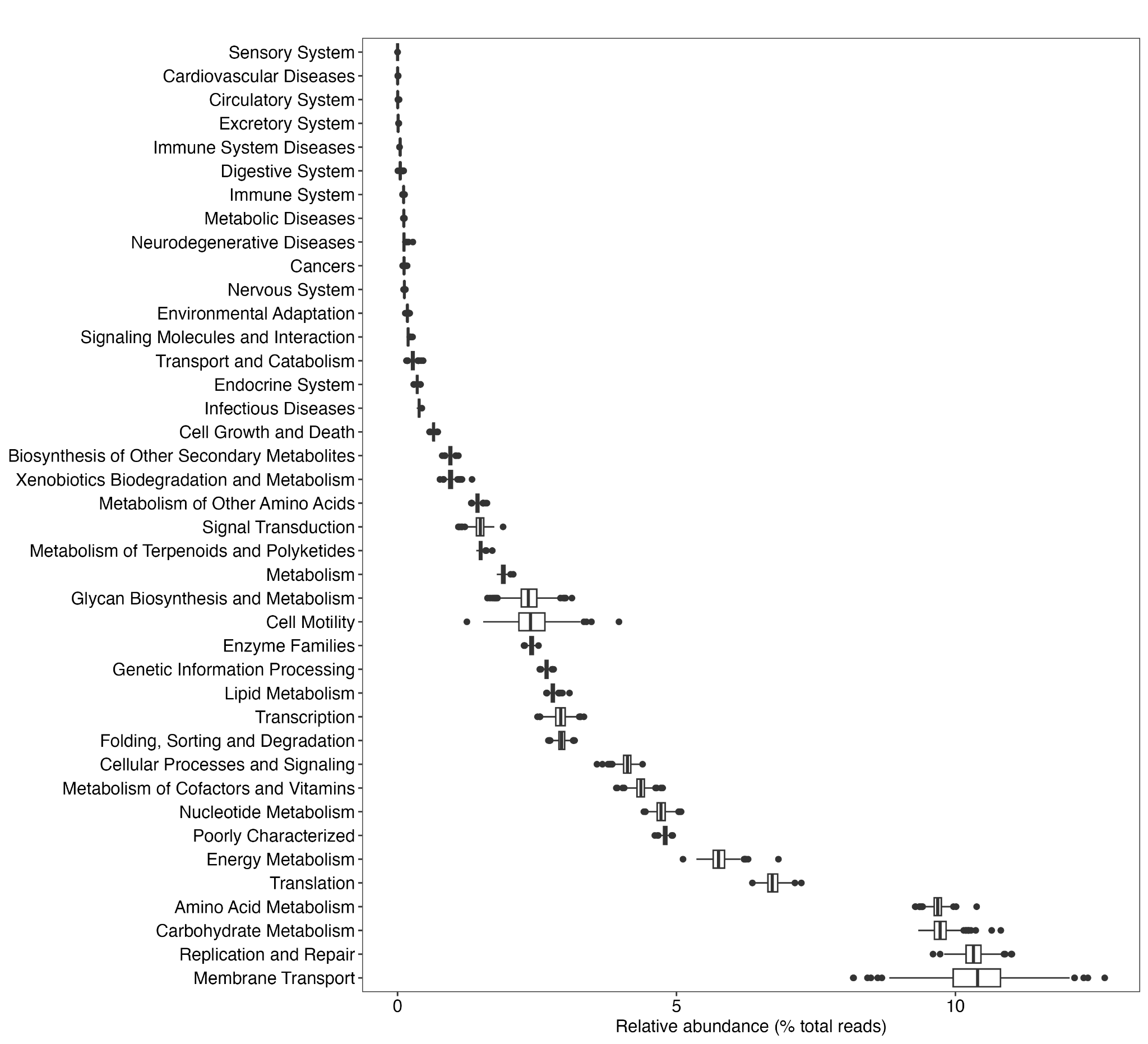
